# Rapid kinetics of H^+^ transport by membrane pyrophosphatase: evidence for a “direct-coupling” mechanism

**DOI:** 10.1101/2024.12.06.627301

**Authors:** Viktor A. Anashkin, Alexander V. Bogachev, Marina V. Serebryakova, Elena G. Zavyalova, Yulia V. Bertsova, Alexander A. Baykov

**Author notes:** Corresponding author (A.A. Baykov). Equal first authors (Viktor A. Anashkin, Alexander V. Bogachev). E-mail addresses* (V.A. Anashkin), (A.V. Bogachev).

## Abstract

Stress resistance-conferring membrane pyrophosphatase (mPPase) found in microbes and plants couples pyrophosphate hydrolysis with H^+^ transport out of the cytoplasm. There are two opposing views on the energy-coupling mechanism in this transporter: the pumping is associated with either pyrophosphate binding to mPPase or the hydrolysis step. We used our recently developed stopped-flow pyranine assay to measure H^+^ transport into mPPase-containing inverted membrane vesicles on the timescale of a single turnover. The vesicles were prepared from *Escherichia coli* overproducing the H^+^-translocating mPPase of *Desulfitobacterium hafniense*. Pyrophosphate induced linear accumulation of H^+^ in the vesicles, without evident lag or burst. In contrast, the binding of three nonhydrolyzable pyrophosphate analogs essentially induced no H^+^ accumulation. These findings are inconsistent with the “pumping-before-hydrolysis” model of mPPase functioning and support the alternative model positing the hydrolysis reaction as the source of the transported H^+^ ions. mPPase is thus a first “directly-coupled” proton pump.

## 1. Introduction

Plants, protists, and many prokaryotes contain membrane-bound pyrophosphatase (mPPase; EC 7.1.3.1, formerly 3.6.1.1), a relict cation pump that couples PP_i_ hydrolysis and synthesis to H^+^ and/or Na^+^ transport across membranes [1–7] and confers stress resistance to host organisms [8–10]. mPPase is functionally similar to F- and V-type ATPases but has a much simpler structure, being a homodimer of 66–89-kDa polypeptides, each folded into 15–17 transmembrane α-helices [11–13] (Figure 1). Each polypeptide forms a funnel-like structure with a hydrolytic center in the cytoplasmic part. The center contains protein ligands for pyrophosphate, four or five Mg^2+^ ions, and nucleophilic water molecule. A gated exit channel protrudes from the hydrolytic center to the vacuolar lumen (in plant vacuoles) or periplasmic space.

**Figure 1.**
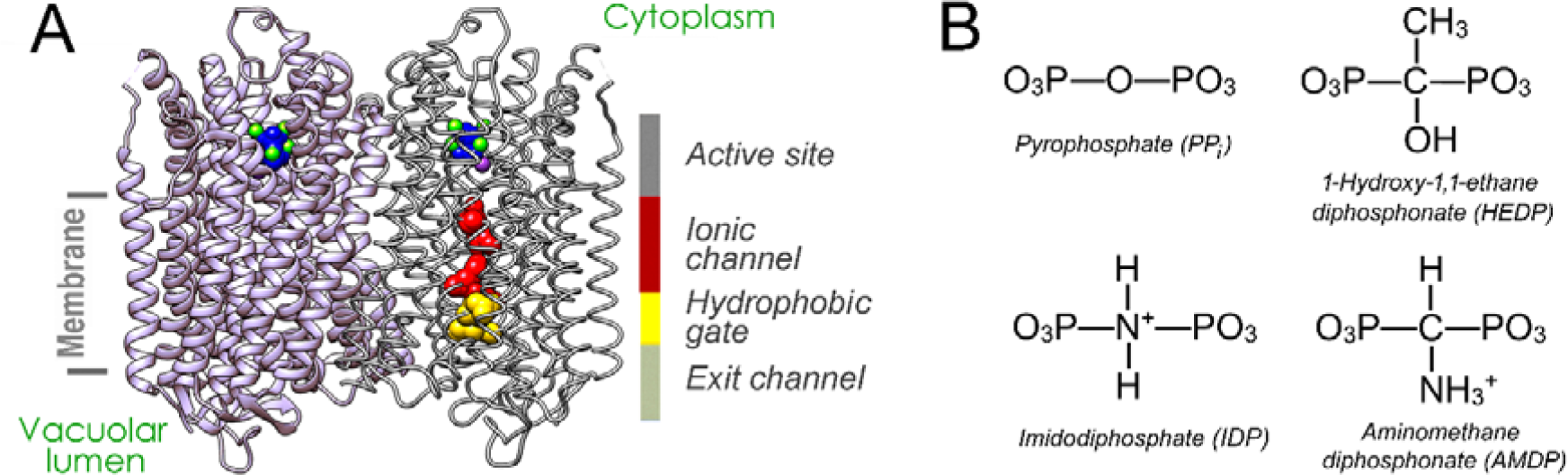
The structures of mPPase and substrate analogs used in this study. (A) Crystal structure of *Vigna radiata* H^+^-translocating pyrophosphatase dimer (PDB code: 4A01) [11]. Helical elements appear as spiral ribbons in the left subunit and as parts of the skeletal structure in the right subunit. The active-site bound substrate analog imidodiphosphate (IDP, blue), five Mg^2+^ ions green), a K^+^ ion (violet), and the residues forming the ionic channel (red) and hydrophobic gate (yellow) are shown in a space □ filling mode in the right subunit. (B) Chemical structures of PP_i_ and its analogs. Panel A was reproduced from Baykov et al. [25] under the terms of the Creative Commons CC BY license.

The close proximity of the hydrolytic center and exit channel in mPPase allows for a unique energy coupling mechanism that is distinct from that of any other H^+^ pump. In this mechanism, generally referred to as “direct coupling” [14], the transported proton is generated from one of the reactants (nucleophilic water molecule) in the coupled chemical reaction [5, 11]. Consistent with the “direct-coupling” mechanism, most studies have indicated a 1:1 H^+^/PP_i_ stoichiometry for mPPase [15–18]. The waterborne proton may also be involved in Na^+^ transport by Na^+^-specific mPPases via a “billiard” mechanism [5, 19]. Notably, all known proton pumps operate via “indirect-coupling” mechanism, in which the transported proton comes from the medium.

The order in which the hydrolysis and H^+^ transport reactions occur in the mPPase coupling mechanism remains, however, a major controversy. Based on the first crystal structure of mPPase [11], Lin et al. considered hydrolysis ahead of transport in their original formulation of the mechanism. The proton released during hydrolysis creates high local acidity, generating a proton current through short proton wires formed by water molecules and polar amino acid residues [11]. In contrast, the “pumping-before-hydrolysis” model propagated by Goldman et al. [12,20,21] assumes that ion pumping is triggered by substrate binding, which induces the closure of the active site and transport-promoting helical rearrangements. In this model, the transported ion may originate from the medium, and the subsequent hydrolysis event simply prepares the transporter for the next transport cycle.

Support for the pumping-before-hydrolysis model of H^+^ transport was sought in experiments with nonhydrolyzable PP_i_ analogs containing N or C atom instead of O in the bridge position [20,21]. The Goldman’s model predicted that their binding to mPPase would cause a single transport event. The compounds tested included imidodiphosphate (IDP), whose crystal structure [22] and binding affinity [23] are very similar to those of PP_i_ and 1-hydroxy-1,1-ethane diphosphonate (HEDP) (Figure 1B). H^+^ transport was monitored by measuring the charge current through the mPPase-loaded liposome membrane. A measurable current was indeed detected upon bringing the liposomes into contact with the IDP-containing solution [20,21]. However, this seemingly positive result raises serious doubts. First, HEDP produced no current [20,21] despite being able to bind [23]. Second, the IDP-induced current was much lower than the current produced by PP_i_ within sufficient time, as was noted later [24,25], for only one hydrolysis turnover. Notably, the dead time of the instrument (approximately 50 ms) correspondЫ to approximately 1 turnover of PP_i_ hydrolysis. Finally, the method measures the movements of any charge (not necessarily proton) in the direction perpendicular to the membrane plane. Thus, the signal might have arisen, for example, from the repositioning of charged or polar amino acid residues in the transporter or additional Mg^2+^ binding to mPPase active center. Both effects are known to be associated with PP_i_ and IDP binding [13,20,26].

In the present study, we addressed this issue by measuring rapid kinetics of proton transfer in membrane vesicles produced from the *Escherichia coli* cells overproducing the H^+^-translocating mPPase of *Desulfitobacterium hafniense*. Importantly, the transport assay used in our study employed the pH indicator dye pyranine and was thus absolutely specific for protons. The effects of the PP_i_ analogs measured using this assay differ from those previously measured by electrometry, supporting the pumping-after-hydrolysis model of H^+^ transport.

## 2. Materials and Methods

### 2.1. Materials

Pyranine was obtained from Eastman Kodak (Rochester, NY, USA); Tris (Trizma base) and valinomycin were obtained from Sigma-Aldrich Co (St Lous, MO, USA); nigericin was obtained from Calbiochem (San Diego, CA, USA), Mops was an Amresco (Solon, OH, USA) product; magnesium sulfate hexahydrate and potassium chloride were obtained from Merck (Rahway, NJ, USA). Imidodiphosphate was prepared as described [27,28]. The potassium salt of aminomethylene diphosphonate (AMDP) was obtained by neutralizing its free acid (a gift of Dr. S. Komissarenko) with KOH.

### Production of mPPase in Escherichia coli and isolation of inverted membrane vesicles (IMV)

The plasmid pET-36b (Novagen, Merck KGaA, Darmstadt, Germany) containing the *Desulfitobacterium hafniense* mPPase (*Dh*-mPPase) gene was expressed in *E. coli* cells as described elsewhere [29], with some modifications to increase *Dh*-mPPase yield. Specifically, different host cells [C43(DE3)] were used, and their expression was carried out in LB broth medium with Novagen Overnight Express Autoinduction System 1. *E. coli* cells were harvested via centrifugation (10,000 g, 10 min) and washed with medium containing 85 mM NaCl, 5 mM MgSO_4_, and 10 mM Tris-HCl, pH 7.5 and further with medium containing 85 mM NaCl and 5 mM MgSO_4_. The sediment was suspended in medium A (10 mM Mops-KOH, pH 7.5, 100 mM KCl, 5 mM MgSO_4_) containing 5 mM pyranine and traces of benzonase. The mixture was passed once through a French press at 16,000 psi, the unbroken cells and cell debris were removed by centrifugation at 27,500 *g* (5 min), and the IMV were sedimented at 150,000 *g* (50 min). The vesicles were washed with medium B (20 mM Mops-KOH, pH 7.5, 100 mM KCl, 5 mM MgSO_4_) via two cycles of resuspension/centrifugation (150,000 *g*, 40 min) and immediately used for transport and hydrolytic activity measurements. IMV were quantitated in terms of their protein content using a bicinchoninic acid method [30] with bovine serum albumin as a standard.

### 2.3. Assay of the transport activity

Stopped-flow measurements of the fluorescence time courses upon PP_i_ or PP_i_ analog addition to the IMV containing entrapped pyranine were performed as described previously [19], with some modifications. A rapid kinetics apparatus (BioLogic Science Instruments, Seyssinet-Pariset, France) consisting of an SFM-3000/S stopped-flow mixer and MOS-200 optical system with a 3-µL cuvette was used. The excitation wavelength was set to 458 nm (monochromator slit width of 4 nm), and emission was detected at wavelengths >495 nm using a SCHOTT GG495 filter. Equal volumes of the IMV suspension, containing 4 µM valinomycin, and solution of PP_i_ or its analogs, both in buffer B, were mixed at 25 □ at a flow rate of 1.3 mL/s (dead-time of 1.5 ms), and fluorescence was monitored for 2 s. The resolution was 0.2 ms in the 0-200 ms interval and 10 ms thereafter. Data from approximately 100 shots were averaged for each curve. The initial rates of the transport reaction (*v*_0_) were determined by fitting the data collected in the 2– 100 ms time (*t*) range to a single exponent (Equation 1) [19], where *F* is the fluorescence, *k* is the rate constant, and the ratio *v*_0_/*k* is the amplitude of the signal change.

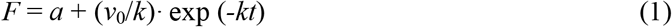

### 2.4. Assay of hydrolytic activity

The rates of PP_i_ hydrolysis were measured at 25 □ using a continuous P_i_ assay, as described previously [19]. The reaction mixture of 20 ml volume contained 100 mM Mops-KOH buffer, pH 7.2, 5 mM MgSO_4_, 25 mM K_2_SO_4_, and IMV (2–8 µg protein/mL). The reaction was started by the addition of 150 µM PP_i_ (tetrasodium salt), and P_i_ liberation was monitored for 3–4 min.

### 2.5. SDS-PAGE

Gel electrophoresis was performed in 4-16% gradient polyacrylamide Tris-glycine gels with 0.1% SDS [31]. The boiling step was omitted because it often causes partial loss of hydrophobic proteins [32]. IMV load was 50 µg protein per lane, and the prestained 10–250 kDa protein ladder PageRuler Plus (Thermo Fisher Scientific Baltics, Vilnius, Lithuania) was run in parallel. The gels were stained with InstantBlue Coomassie (Abcam, Cambridge, UK) and quantified using Bio-Rad Bio-Rad (Hercules, CA, USA) ChemiDoc MP imaging system and ImageLab (ver. 5.1) software.

### 2.6. Mass spectrometry

MALDI-TOF MS analysis of trypsin-produced peptides was performed using an UltrafleXtreme MALDI-TOF-TOF mass spectrometer (Bruker Daltonik, Bremen, Germany) as described elsewhere [33]. Protein identification was performed by MS+MS/MS ion search, using Mascot software version 2.3.02 (Matrix Science, USA) through the Home Protein Database.

## 3. Results and Discussion

### 3.1. Production and quantification of Dh-mPPase in IMV

To reliably detect H^+^ translocation within a single catalytic cycle, high mPPase concentration must be maintained in the assay. *Dh*-mPPase was efficiently produced in *E. coli* C43(DE3) cells transformed with a plasmid containing *Dh*-mPPase gene under T7 promoter, as indicated by activity measurements and SDS-PAGE analysis of the membrane fraction of cell homogenate. Two IMV types were analyzed: one obtained from cells transformed with the above plasmid and the other obtained from the cells transformed with the plasmid lacking the *Dh*-mPPase gene (“empty” pET-36b plasmid). The activities of these IMV preparations in PP_i_ hydrolysis were typically 1.0–1.1 and < 0.01 µmol/min per 1 mg protein, respectively.

In accordance with these findings, SDS-PAGE analysis of the *Dh*-mPPase-containing IMV revealed two additional protein bands with the apparent masses of 56 and 110 kDa in comparison with the IMV produced from the cells lacking the *Dh*-mPPase gene (Figure 2). A wide-range gradient gel was used to allow separation of the whole set of membrane proteins.

**Figure 2.**
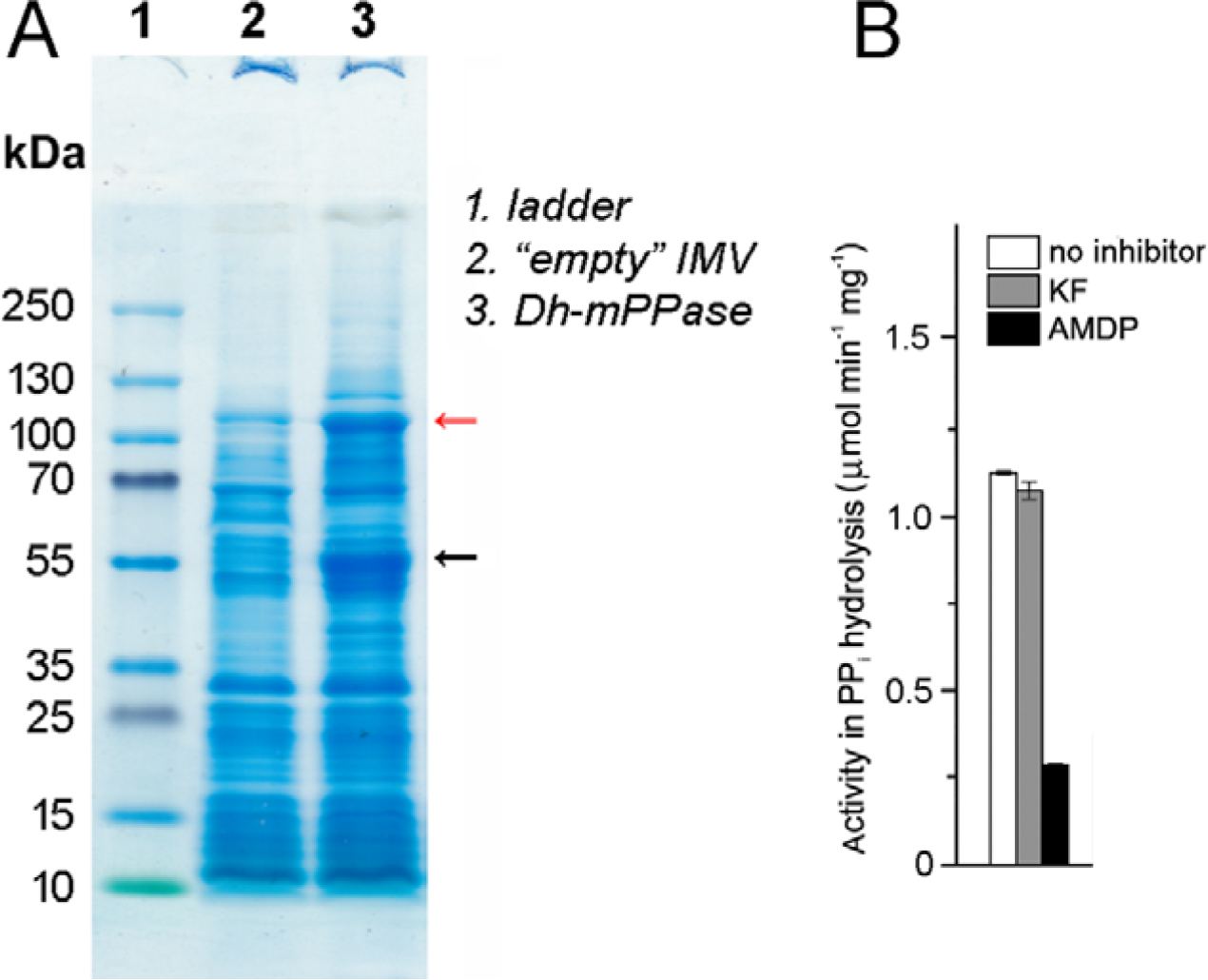
Production of *Dh*-mPPase in *E. coli*. (A) Coomassie staining of *E. coli* membrane proteins separated by SDS–PAGE. The putative mPPase bands are indicated by arrows. The leftmost lane shows reference molecular mass markers. (B) Activities of IMV in PP_i_ hydrolysis. The inhibitors potassium fluoride (1 mM) and AMDP (20 µM) were present where indicated.

Standard MALDI-TOF MS- and MS/MS-analyses were performed to identify the proteins present in the two protein bands. The results indicated that the 110-kDa band contains a E1 subunit of *E. coli* pyruvate dehydrogenase complex, whereas the 56-kDa band mainly belongs to *Dh*-mPPase but contains approximately one third by mass *E. coli* dihydrolipoyl dehydrogenase and traces of ATP synthase subunit α (Supplement). The theoretical mass of *Dh*-mPPase subunit is 69 kDa, but, because of its high hydrophobicity and like all with other mPPases [29], it migrates faster in PAGE than a soluble protein of the same mass.

The contribution of the *Dh*-mPPase band to the total intensity of the stained bands on the scanned gel was estimated with the program ImageLab and found to be 7–8 %. This value roughly corresponds to the contribution of *Dh*-mPPase to the total IMV protein. Based on the above percentage and the specific activity of IMV in PP_i_ hydrolysis and taking into account that mPPase contains only one functional active site per dimer [34], the turnover number of individual *Dh*-mPPase can be calculated as 32 s^-1^. This value exceeds the range of 11.5–18.5 s^-1^ reported for other H^+^-translocating mPPases [35–37].

### 3.2. Fast kinetics of transmembrane proton transport by Dh-mPPase

*Dh*-mPPase-catalyzed acidification of the IMV lumen was followed by measuring a decrease in the fluorescence of the pH indicator pyranine encapsulated in IMV [19,38]. Pyranine fluoresces only in its deprotonated form (*pK*_*a*_ = 7.2) [38] and exibits a very fast response rate [39]. The IMV containing the encapsulated pyranine was rapidly mixed with PP_i_ solution in a stopped-flow instrument with a mixing dead-time of 1.5 ms. Based on the determined *Dh*-mPPase content (∼7.5 % of IMV protein, corresponding to ∼0.55 nmol *Dh*-mPPase per 1 mg protein) and the transport stoichiometry of 1 H^+^/PP_i_, one would expect an ∼7.5 mmol/L increase in the H^+^ content inside IMV upon a single synchronized turnover of IMV *Dh*-mPPase. This estimate is high because of the very low specific volume of IMV (∼0.07 µl per 1 mg protein [40]). To further boost the fluorescence signal associated with H^+^ transport into IMV, the pyranine concentration inside IMV was increased from 1 to 5 mM and the assay pH was increased from 7.2 (equal to pyranine p*K*_a_) to 7.5 in comparison to the previous publication [19]. Theoretically, our assay provides the possibility of monitoring H^+^ transport even at a high buffering capacity of the medium inside IMV.

The red curve in Figure 3A shows the time course of pyranine fluorescence over 2 s after mixing *Dh*-mPPase-containing IMV with 75 µM PP_i_. The reaction medium contained the K^+^-ionophore valinomycin to prevent the limitation of the transport reaction by the built transmembrane electrical potential difference. A rapid decay with *t*_1/2_ of approximately 140 ms to a constant level was observed. The amplitude of the signal In Figure 3A was apparently limited by the amount of encapsulated pyranine, which was finally fully converted into a protonated form. No transport was observed without PP_i_ (Figures 3A and 3B, black curves) or with IMV produced from the recombinant *E. coli* strain lacking the *Dh*-mPPase gene (data not shown). The K^+^/H^+^ exchanger nigericin inhibited H^+^ transport (Figure 3C). These findings confirmed that the pyranine signal is indicative of the H^+^ transport reaction catalyzed by *Dh*-mPPase.

**Figure 3.**
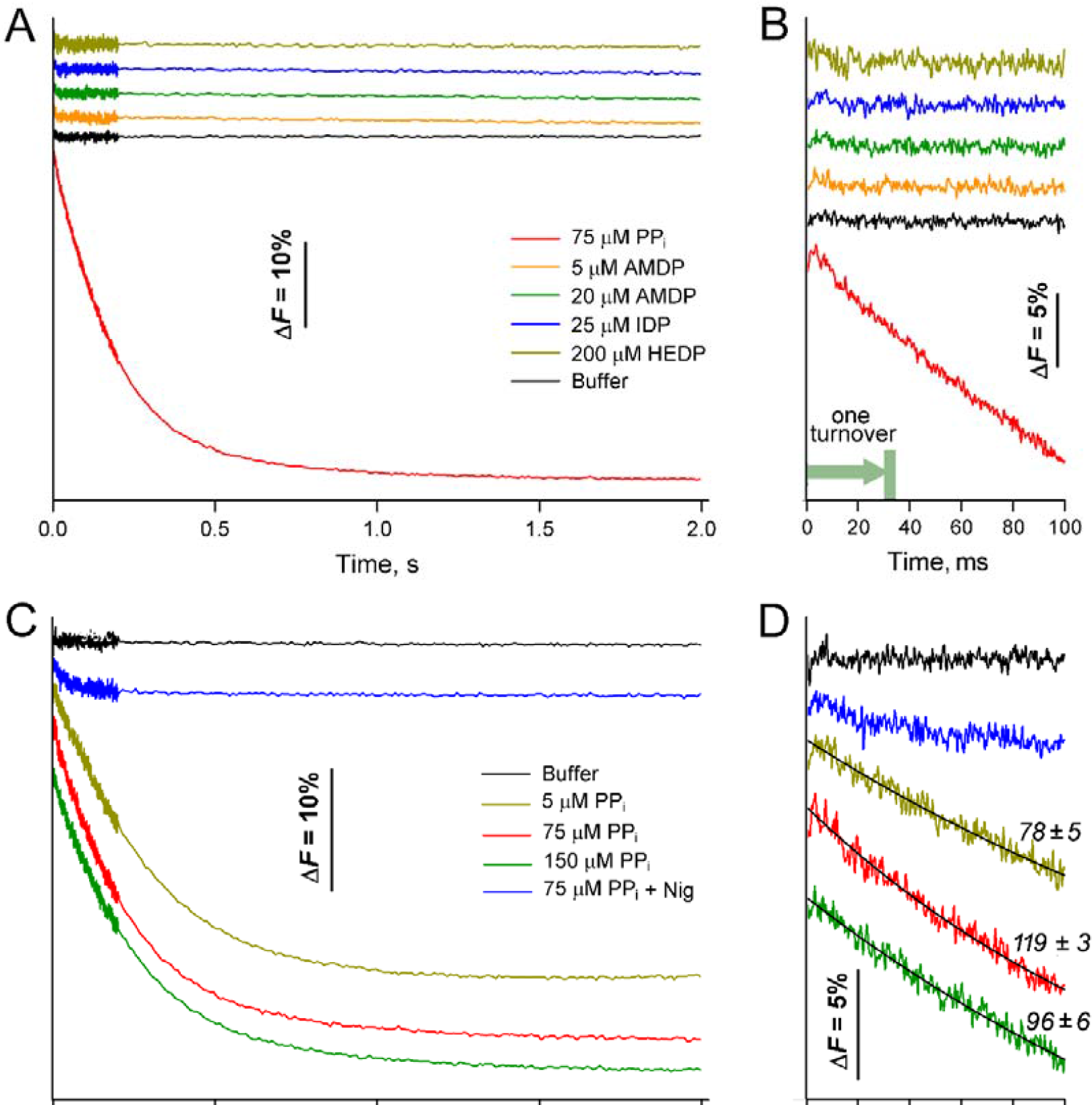
Typical stopped-flow traces of pyranine fluorescence recorded after combining the membrane vesicles harboring *D. hafniense* H^+^-translocating mPPase with PP_i_ or its analog. Vesicle suspension (0.30–0.36 mg protein/mL, 20 mM Mops-KOH, pH 7.5, 100 mM KCl, 5 mM MgSO_4_, and 4 µM valinomycin) was preincubated for 10 min and mixed with an equal volume of PP_i_ or PP_i_ analog solution in the same medium without any ionophore. Two different IMV batches were used for panels A and C. (A) Mixing with fixed concentrations of PP_i_ or PP_i_ analogs (detailed in panel label). The curves were shifted vertically for clarity. (B) Enlarged view of the kinetic curves in panel A in the 2–100 ms range. (C) Effects of varying PP_i_ concentration and adding nigericin (4 µM in the IMV suspension). (D) Enlarged view of the kinetic curves in panel C in the 2–100 ms range. The black curves show the best fits of Equation 1 with the *v*_0_ values (in % fluorescence per second) shown on the curves.

Based on the turnover number determined for *Dh*-mPPase, the change in the fluorescence signal by approximately 5 % during the first 30 ms of the transport reaction is associated with one synchronized catalytic cycle. Importantly, the fluorescence time-course was virtually linear in this time interval and demonstrated no initial lag or burst (Figure 3B). Thus, the transport assay is sensitive enough to reliably detect the H^+^ transport resulting from a single turnover of Dh-mPPase (i.e. within a 30 ms period) (Figure 3B).

We therefore used this assay to directly test the “pumping-before-hydrolysis” model of mPPase functioning, which predicts a single act of H^+^ transport upon non-convertible PP_i_ analog binding to *Dh*-mPPase. The analogs contained a C or N atom instead of an O atom in the bridge position and included AMDP, IDP, and HEDP (Figure 1B). Like pyrophosphate, they bind with largely different affinities to two active sites of the enzyme diner, demonstrating strong negative cooperativity [34]. The respective *K*_i1_ values corresponding to inhibitor binding to one active site are 0.40, 4.2, and 27 µM. For comparison, the Michaelis constant (*K*_m_) is 2.3–2.6 µM in terms of Mg_2_PP_i_ (the “true” substrate of mPPase) [34], or 3.4–3.9 µM in terms of total PP_i_, as used in the present work. At the analog concentrations used, which were limited by AMDP and IDP solubilities, the first site was thus expected to be 83–98 % saturated. The occupancies of the second site by IDP and HEDP were <20%, but AMDP (*K*_i2_ of 12.5 µM) was expected to occupy 61 % of the second sites [34] at its 20 µM concentration. At 5 µM AMDP, occupancies of 93 and 28 % were expected for the two sites.

As Figures 3A and 3B highlight, binding of neither substrate analog induced an observable change in pyranine fluorescence on the timescale of milliseconds or seconds. This finding disagrees with the “pumping-before-hydrolysis” model and suggests that the limited charge transfer observed upon IDP binding to H^+^-translocating mPPase in Goldman’s lab [20,21] is not associated with H^+^ transport. As our calculations indicated that AMDP can occupy both active sites in the enzyme dimer, the inability of AMDP to effect H^+^ transport also rules out a possible model linking the membrane transport with the substrate occupancy of the second active site.

The H^+^ transport reaction was also assayed at two different PP_i_ concentrations (Figure 3C), and the initial rates of the associated fluorescence change *v*_0_ were estimated using Equation 1 and presented in Figure 3D. The rate was maximal with 75 µM PP_i_ and decreased when PP_i_ concentration was either decreased to 5 µM or increased to 150 µM, in accordance with the effects on the hydrolytic activity [34]. *Dh*-mPPase inhibition by excess substrate is explained by its binding to the second active site in the dimer with a concomitant decrease in *k*_cat_ value for both sites [34]. Only a modest dependence of the initial rate of H^+^ transport on PP_i_ concentration indicated that transporter turnover is not limited by substrate binding step—the latter is a second-order reaction with its rate proportional to substrate concentration. These data agree with recent findings showing that the catalytic cycle of Na^+^-transporting mPPase is limited by the hydrolysis step, rather than pyrophosphate binding or phosphate release steps [41]. Thus, our new data support the notion that pyrophosphate hydrolysis is the coupling step in mPPase-catalyzed proton transfer.

## 4. Conclusions

The success in *Dh*-mPPase overproduction in *E. coli*, small inner volume of IMV, and synchronous starting of the hydrolysis reaction allowed resolution of a single turnover of the H^+^ transport reaction. The use of nonconvertible tightly binding PP_i_ analogs provided strong experimental support to the theory that proton translocation by mPPase is coupled to the chemical step rather that substrate binding. Theoretically, the absence of a transport signal on the curves measured with the PP_i_ analogs could have been a consequence of very fast pumping during the 1.5-ms deadtime of the stopped-flow instrument. Notably, a slow “alkaline” response of pyranine fluorescence with an amplitude of 8 % fluorescence signal on the time scale of seconds, would be expected in this case, caused by relaxation of the ΔpH formed in the single turnover of the proton transporter. However, this relaxation was not observed, ruling out this hypothesis.

## CRediT authorship contribution statement

**Viktor A. Anashkin:** Formal analysis, Investigation, Methodology, Validation, Visualization, Writing – original draft. **Alexander V. Bogachev:** Conceptualization, Investigation, Methodology, Validation, Writing – original draft. **Marina V. Serebryakova:** Resources, Methodology, Formal analysis. **Elena G. Zavyalovs:** Resources, Methodology. **Yulia V. Bertsova:** Investigation, Validation. **Alexander A. Baykov:** Visualization, Funding Acquisition, Writing – review and editing.

## Funding sources

This study was funded by the Russian Science Foundation (research project 23-24-00115).

## Declaration of competing interest

The authors declare no competing interest.

## Acknowledgments

The stopped-flow apparatus of the MSU Shared Research Equipment Center ‘Subdiffractional Microscopy and Spectroscopy’ and the MALDI MS facility became available in the frameworks of the Equipment Renovation Program of the National Project ‘Science’ and the Moscow State University Development Program PNG 5.13, respectively.

## Appendix A. Supplementary data

Supplementary data to this article can be found online at https://

## Notes

### Competing Interest Statement

The authors have declared no competing interest.

### Summary of Updates

Missing CRediT and funding information added.

